# When Tagging Frequency Matters to Attention: Effects on SSVEPs, ERPs, and Cognitive Processing

**DOI:** 10.64898/2026.03.30.715193

**Authors:** Jihan Yang, Olivia Carter, Mohit N Shivdasani, David B. Grayden, Rob Hester, Ayla Barutchu

## Abstract

Selective attention enables the prioritization of task-relevant information while managing distractors, and steady-state visual evoked potentials (SSVEPs) are widely used to track this process by tagging different visual objects at distinct flicker frequencies. However, whether the choice of tagging frequency itself influences other neural and cognitive measures remains unclear. Here, 27 participants performed detection and 1-back working memory tasks while a central target and peripheral distractors flickered at either 8.6 Hz or 12 Hz. The working memory task produced slower responses, more errors, and greater perceived difficulty than detection. Tagging frequency strongly shaped neural responses, with 8.6 Hz eliciting higher SSVEP signal-to-noise ratios than 12 Hz regardless of stimulus location. Nevertheless, stronger SSVEP responses for centrally attended stimuli were associated with fewer working memory errors and larger early visual ERP responses, while SSVEPs for attended and distractor stimuli were negatively correlated. In addition, the working memory task produced a larger P1-N1 peak-to-peak difference, and tagging frequency altered the timing and amplitude of early ERP effects. Together, these findings show that tagging frequency is not a neutral methodological parameter, but one that shapes both neural indices of attention and their relationship to cognitive performance.

## Introduction

The daily environment is cluttered and dynamic and, more often than not, humans significantly rely on the visual system to selectively attend to particular stimuli and tasks while ignoring surrounding distracting objects and events. Understanding selective visual task-specific functions is important not only for comprehending human behavior but also for developing interventions to improve performance in educational, organizational and clinical settings. Steady-state visual evoked potentials (SSVEPs) have become a cornerstone technique in cognitive neuroscience for investigating concurrent neural processes, as they enable dissociation of neural mechanisms related to multiple simultaneously processed objects and events [1, 2]. This dissociation is often achieved by frequency-tagging multiple visual objects by presenting them at distinct frequencies (e.g., 8.6 Hz and 12 Hz as in the present study) to modulate neural oscillations for different tasks and objects.

SSVEPs demonstrate robust modulation by attention, with enhancements for attended compared to ignored stimuli [3–6]. The presence of distracting objects can reduce the SSVEP magnitude [7], though responses remain greater for attended than for distractor objects [8]. Recent investigations of distractor processing using frequency-tagging paradigms have revealed that learned distractor suppression can modulate SSVEP responses within approximately 180 ms post-stimulus onset [9], and that competitive effects between emotionally arousing distractors and task-relevant stimuli emerge around 450 ms and persist for several seconds [10]. Furthermore, when task demands require divided attention between overt and covert targets, SSVEPs show transient reductions that reflect dynamic shifts in attentional allocation [11]. These findings suggest that attention operates as a limited resource that spreads to distractors despite selectivity toward attended objects.

The spatial distribution of attentional allocation varies depending on task demands. For overlapping objects, unattended features may be suppressed [3]. However, for peripheral distractors, attention appears to spread and upregulate neural activity to task-relevant features rather than exclusively suppressing distractors with task-irrelevant features [12]. Recent evidence using electrophysiological measures in visual search has demonstrated that highly salient distractors may initially capture attention without being suppressed below baseline, with attention subsequently focusing upon targets [13]. This pattern is consistent with center-surround configurations of load-induced enhancement and suppression in the visual field, where high perceptual load attenuates distractor SSVEPs at proximal eccentricities but not at more peripheral locations [14]. Moreover, the relationship between working memory and attentional allocation in SSVEP paradigms has been clarified by recent work showing that SSVEPs can track dynamic reallocation of spatial attention during visual working memory maintenance, with SSVEP coherence decreasing at distractor locations shortly before memory probe comparison [15]. This suggests that memory demands modulate the spatial distribution of attention in ways that can be captured by frequency-tagging.

The relationship between SSVEP responses and tag frequency presents additional complexity. While attention effects on SSVEPs have been shown to be consistent across different frequency tags in simple detection and search tasks [7, 12, 16], SSVEP patterns may vary with tag frequency depending on target object familiarity [17] and the spatial location of stimuli [18]. Critically, the choice of flicker frequency determines which cortical network synchronizes to the stimulus, with distinct networks showing different attention effects [18]. Recent magnetoencephalography studies have revealed that spatial attentional modulation patterns differ systematically across subdivisions of early visual cortex depending on the tagging frequency employed, with frequencies in the 8-15 Hz range providing optimal signal-to-noise ratios [19]. Furthermore, temporal dynamics of SSVEP responses under sustained stimulation exhibit frequency-dependent patterns, with lower frequencies (e.g., 5 Hz) showing progressive amplitude increases while higher frequencies demonstrate initial increases followed by continuous decline [20]. These findings highlight the need to consider tag frequency as a potential moderating variable when interpreting attention-related SSVEP effects, particularly in paradigms that employ multiple tagging frequencies to simultaneously track central and peripheral processing.

Despite these advances, no study has systematically investigated how the task relevance of distractor stimuli affects SSVEPs when central attended targets and peripheral distractors share identical familiar features, with only the task relevance of distractors differing between conditions. Furthermore, how SSVEPs and their tag frequencies relate to task performance and participants’ subjective experiences remains poorly understood. Critically, while SSVEPs index sustained attentional engagement with frequency-tagged stimuli across an entire trial, they do not capture the rapid, early transient neural responses that occur during early stages of visual processing. Early ERP components such as P1 and N1 (70–180 ms) provide complementary, time-resolved measures of perceptual and attentional processing that may reveal how the brain initially allocates resources under different task demands prior to neural entrainment (i.e., the onset of SSVEPs). The present study aimed to address these gaps by comparing SSVEP responses across a simple detection task and a working memory 1-back task (manipulating task relevance of peripheral distractors), using two distinct tagging frequencies (8.6 Hz and 12 Hz) to simultaneously track the processing of centrally attended task relevant stimuli and peripheral distractors. We predicted that SSVEP amplitude would be greater for the working memory task than for the detection task, and that the relative difference in activation between central targets and peripheral distractors would be modulated by both task relevance of distractors and the tag frequency.

## Methods

### Participants

Thirty-four participants were recruited for this study. Four participants were excluded due to recording failure. Two were excluded from all analyses due to high error rates (>30% error) and one excluded for excessive EEG artifacts (< 30 trials after artifact rejection), yielding a final sample of 27 participants (*M* age = 24.59 years, *SEM* = 1.30, age range 18-43 years, 8 males). All participants had normal hearing, normal or corrected-to-normal vision, and no history of neurological or psychiatric disorders. Participants were randomly allocated into two frequency-tag order groups: Group C8 (n = 14, central target presented at 8.6 Hz, peripheral distractors presented at 12 Hz) and Group C12 (n = 13, central target presented at 12 Hz, peripheral distractors presented at 8.6 Hz). All procedures were approved by and strictly adhered to the guidelines of the University of Melbourne Human Research Ethics Committee. The study was conducted in accordance with the Declaration of Helsinki. Written informed consent was obtained from all participants prior to the experiment.

### Experimental design and stimuli

During experiments, participants were seated in a dimly lit electrically shielded room at a viewing distance of 70 cm from a 19-inch CRT monitor (refresh rate: 120 Hz) and asked to fixate centrally on a fixation cross (0.4° visual angle). Stimuli consisted of five white capital letters from the English alphabet (Helvetica font) presented on a grey background: one central target letter and four peripheral distractor letters positioned along the horizontal and vertical axes. Each letter subtended a visual angle of 3.3°. To minimize spatial interference between the SSVEP tags, the distractor letters were positioned at a visual angle of 4.9° relative to the central letter.

To elicit SSVEPs, the central target and the peripheral distractors were flickered at 8.6 Hz and 12 Hz, respectively, or vice versa (counterbalanced across stimulus locations and task conditions). These frequencies were selected to ensure sufficient spectral separation while remaining within the alpha band range commonly used in SSVEP attention studies [7, 8, 18]. Task order was also counterbalanced across participants. For each trial, distractor letters were randomly selected from all possible consonants (excluding letters presented in previous trials). The central target letter was randomly selected on the first trial of each block and subsequently drawn from one of the four distractor stimuli of the previous trial.

Each trial lasted for 4083 ms, followed by an interstimulus interval (ISI) that randomly varied between 1 and 2 s (Figure 1a). Participants performed two tasks using identical stimulus presentation sequences, with only the task instructions differing: a 1-back working memory task and a simple detection task. In both tasks, the target was a single red color change of a letter, lasting 300 ms, which occurred at a random time between 300 ms and 3500 ms post-stimulus onset. This red flash was a one-time color substitution overlaid on the continuously flickering letter; it was not presented at the flicker rate but replaced the white letter with a red letter for the flash duration while the underlying flicker continued. Participants were instructed to strictly ignore peripheral distractors in both tasks.

**Figure 1.**
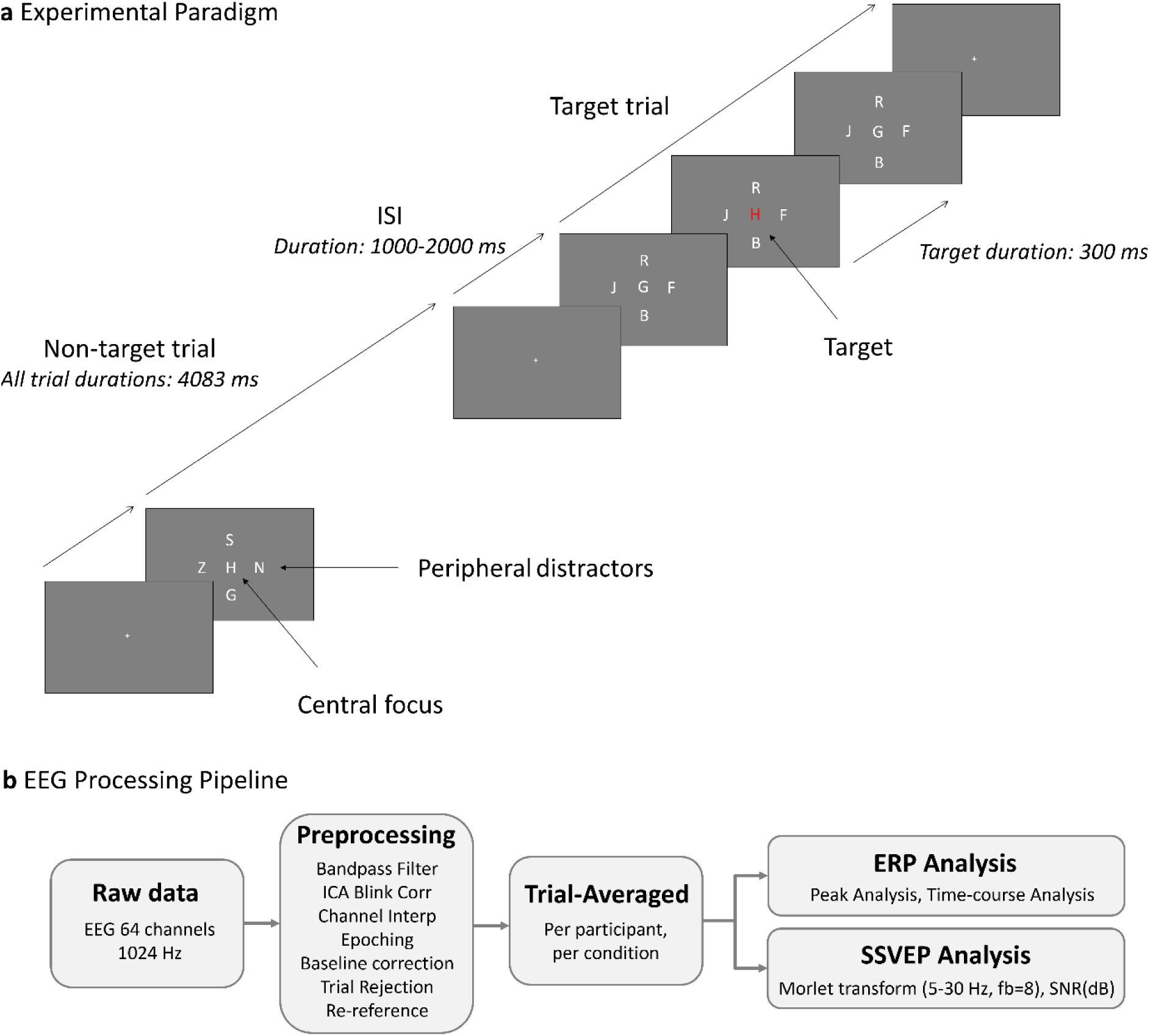
**a.** Schematic of two consecutive trials illustrating the detection and working memory tasks. In the working memory task, only a red letter matching the central letter of the previous trial constituted a valid target (e.g., red ‘H’); in the detection task, any red letter was a valid target regardless of identity. **b.** Overview of the EEG preprocessing and analysis pipeline.

For the working memory task, participants were required to remember the central white letter and press a button if a red matching letter flashed during the subsequent trial (Figure 1a). To ensure continuous attention and prevent participants from merely responding to any color or identity change, three types of catch trials were included (change in color only, change in letter identity only, and change in both non-matching identity and color). This constituted a dual-load 1-back task: encoding the current white letter while actively searching for the red match of the previous trial. This procedure ensured that both the color and the letter identity remained relevant, and attention was maintained throughout the entire trial.

For the detection task, participants ignored letter identity and pressed a button whenever they detected a red color flash. Participants completed short practice runs (<50 trials) followed by two blocks of 100 trials for each task. Each block comprised 70% no-change (invalid) trials, 20% valid target trials, and 10% catch trials (for the 1-back task).

### Subjective measures

At the completion of each task, participants completed five 10 cm visual analogue scales (VAS). The VAS assessed: (1) relaxation level (very relaxed to very stressed), (2) alertness (very tired/sleepy to very alert), (3) concentration ease (very easy to very difficult), (4) the perceived difficulty of the task (very easy to very difficult), and (5) exerted effort (very little effort to very much effort).

### EEG recording and preprocessing

EEG data were recorded from 64 scalp electrodes using a BioSemi ActiveTwo system at a 1024 Hz sampling rate. The BioSemi system uses a Common Mode Sense (CMS) and Driven Right Leg (DRL) reference scheme during online recording. Two additional electrodes were placed over the left and right mastoids for recording purposes but were not used as references or included in subsequent analyses. Horizontal and vertical electrooculography (EOG) channels were collected to monitor eye movements and blinks. A photodiode (recorded via the BioSemi Erg1 external channel) was attached to the corner of the CRT monitor to verify the timing of the visual flicker.

EEG preprocessing was performed using EEGLAB [21] and MATLAB (Figure 1b). Data were bandpass filtered between 1-45 Hz to remove low-frequency drift and high-frequency noise. Only behaviorally correct non-target trials were retained in the EEG analysis (i.e., catch trials and trials that required a motor response were excluded). Additionally, a photodiode was used to record and verify flicker validity using power spectral analysis. A criterion was set whereby trials with insufficient photodiode flicker-related power at the target frequency (absolute spectral power < 1.56 × 10⁹), determined via power spectral analysis, would be excluded. This threshold was used to identify trials in which the visual flicker was not reliably presented (e.g., due to stimulus rate errors). No trials met this criterion.

Independent Component Analysis (ICA) was performed using the Picard algorithm [22]. Eye-movement related components were identified using ICLabel [23] and removed if their classification probability exceeded 0.8. Noisy channels were identified using a dual criterion: channels whose mean power spectral density deviated by more than ±6 SD from the grand mean across all channels, and whose average correlation with the four nearest neighboring channels fell below 0.6. Both criteria had to be met simultaneously for a channel to be flagged. The automated detection was spot-checked on a subset of participants to confirm validity. Flagged channels were spherically interpolated rather than discarded to preserve spatial information.

Continuous EEG data were segmented into epochs from −600 ms to 4600 ms relative to each trial onset. Baseline correction was applied separately for ERP and SSVEP analyses: a −200 to 0 ms window was used for ERP analysis, while a −600 to −200 ms window was used for SSVEP analysis to avoid edge effects from wavelet convolution. The first trial of all datasets was removed due to the nature of the 1-back task. For each epoch, artifact rejection was applied in two steps: (1) epochs exceeding ±100 µV in any channel were excluded; (2) epochs meeting either of the following two criteria: (a) more than 3% of sample points deviating by ±2.4 SD from the mean at each time point across all trials for each channel, or (b) peak power in the 30–45 Hz range exceeding the mean +3.5 SD, were also excluded. Clean data were then re-referenced to the average of remaining scalp EEG channels (mastoid, EOG, and photodiode channels were excluded in earlier preprocessing steps). Datasets with fewer than 30 clean trials per condition were excluded from analysis (yielding one participant exclusion).

Across the 27 retained participants, a total of eight channels were interpolated: six in the detection task (Fp1, T8, F8, Oz, AF8, T7) and two in the memory task (Iz, O1). For one participant, interpolated channels included Oz and O1, both within the parieto-occipital ROI; given the high electrode density in this region and the use of six-electrode ROI averaging, the impact on SSVEP and ERP measures was minimal. No channels within the frontal ROI required interpolation.

### Data analysis

#### Behavioural data analysis

Reaction times (RTs) greater than 100 ms and within ±3 SD of each participant’s mean were classified as valid responses for target trials. Only participants with error rates below 30% across all stimulus types were included in the analysis. Accuracy was high across stimulus types; therefore, only overall error rates were analysed. Because the assumption of normality was violated for the detection task, group differences for each task were examined using independent-samples Mann–Whitney U tests. Within-group task differences were analysed using related-samples Wilcoxon signed-rank tests.

Mean reaction time (MRT) for correct responses were analyzed using a 2 (Task: Detection vs. Working Memory) × 2 (Group: C8 vs. C12) mixed ANOVA, with Task as a within-subject factor and Group as a between-subject factor. Subjective VAS ratings were analyzed using the same ANOVA design.

#### Event-Related Potentials (ERPs) analysis

Only correct ‘invalid’ trials were included in the ERP and SSVEP analyses (note that during these trials, participants continuously searched for a valid target, as the target could have appeared at any time point throughout the trial). For each individual and condition, clean epochs were averaged to compute ERPs. Statistical analyses of ERPs were restricted to the first 600 ms post-stimulus onset (To visualize the entire ERP −200 – 4600 ms, see Supplementary Information A).

We focused on early visual ERP peak components at Oz and POz that were unlikely to be attenuated by SSVEPs: P1 (70–130 ms, positive peak), N1 (90–190 ms, negative peak), and P2 (150–220 ms, positive peak). For each participant, peak amplitudes and latencies were extracted using MATLAB’s ‘findpeaks’ function with a minimum prominence threshold of 0.01 μV. To avoid noise-related spurious peaks, the peak with the highest prominence within the defined window was selected. If no peak met the criterion, the maximum/minimum value within the window was used as a fallback. Component order was strictly validated to ensure P1 preceded N1 and N1 preceded P2 in latency. In cases where temporal order was violated (e.g., the detected N1 latency fell before the detected P1 latency), the two disordered components were assigned a shared amplitude value, computed as the mean of their individually detected amplitudes, to avoid artifactual reversals in the peak measures. Peak measures were analyzed using 2 (Task) × 2 (Group) mixed ANOVAs in SPSS, with Task as a within-subject factor and Group as a between-subject factor.

We also ran time-course analyses along the ERP. For each ERP sample, point-by-point t-tests were conducted across the post-stimulus epoch (0-600 ms) at electrode Oz and POz. Between-group comparisons (Group C8 vs. Group C12 within each task) used independent-samples t-tests, while within-subject comparisons (Detection vs. Memory within each group) used paired-samples t-tests. To control for multiple comparisons while maintaining sensitivity to sustained effects, a temporal clustering criterion based on the Guthrie and Buchwald (1991) framework was applied [24]. Given our high sampling rate (1024 Hz), significance was defined as a minimum of 20 consecutive milliseconds (21 consecutive data points) reaching significance at α = 0.05.

#### SSVEP Time-Frequency analysis

For each participant and condition, trial-averaged ERPs were decomposed into time–frequency representations from 5 to 30 Hz in 0.2 Hz steps using complex Morlet wavelets (bandwidth parameter fb = 8) with a length of ±3 standard deviations around each frequency. To enable meaningful comparison across different frequencies despite the 1/f characteristics of the EEG power spectrum, the signal-to-noise ratio (SNR) was computed at each tagging frequency (8.6 Hz and 12 Hz) using the formula [25]:

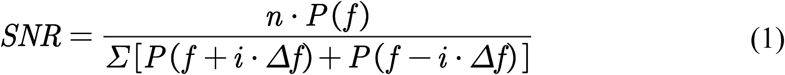

Where, i = 1 to n/2, P(f) is the power at the target frequency, n = 6, and Δf = 0.5 Hz. SNR values were converted to decibels (dB) using

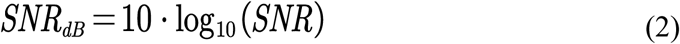

SNR(dB) was then averaged over the 1000-3500 ms time window to avoid onset and offset transients, and across a parieto-occipital region of interest (ROI) comprising six electrodes (PO3, POz, PO4, O1, Oz, O2). Working memory has been associated with SSVEP modulations at frontal sites [26–28]. To examine whether task-related modulations were present at frontal electrodes, a supplementary analysis was conducted using a frontal ROI (Fz, F3, F4), consistent with electrode sites reported in these prior studies [26–28].

SNR(dB) values were analyzed using a three-way mixed ANOVA with Task (Detection vs. Memory) and Location (Central vs. Peripheral) as within-subject factors and Group (8.6 Hz vs. 12 Hz central tagging frequency) as a between-subject factor. The dependent variable was the mean SNR(dB) across the parieto-occipital ROI for each participant under each experimental condition. Unless otherwise specified, significance was evaluated at α = .05. Sidak corrections were applied for pairwise comparisons of main effects, while Bonferroni corrections were used for simple effects analysis of interactions.

## Results

### Behavioural results

Error rates did not differ between Group C8 and Group C12 for the detection (*z* = 1.53, *p* > .05) and memory tasks (*z* = 1.24, *p* > .05). Error rates were significantly higher in the memory than the detection task for both Group C8 (*z* = 2.61, *p* < .01) and Group C12 (*z* = 3.11, *p* < .01). Response times (RTs) were also significantly slower for the working memory task than the detection task, *F*(1, 25) = 51.39, *p* < .001, *η²* = .67. There were no other significant main or interaction effects for RTs.

Participants’ subjective reports on the visual analogue scales for relaxation and alertness, did not differ significantly between tasks. However, participants rated the working memory task to be significantly more difficult, *F*(1, 25) = 35.97, *p* < .001, *η²* = .59, and effortful, *F*(1, 25) = 15.43, *p* < .001, *η²* = .38. The only difference in subjective measures between the tag frequency groups was for the subjective measure of concentration, where participants in Group C8 (central tag 8.6 Hz and peripheral distractor tag 12 Hz) rated the task as marginally easier to concentrate on than Group C12, though this difference was only borderline significant, *F*(1, 25) = 4.11, *p* = .05, *η²* = .14, which may reflect a Type II error as observed power = .50.

**Table 1.**
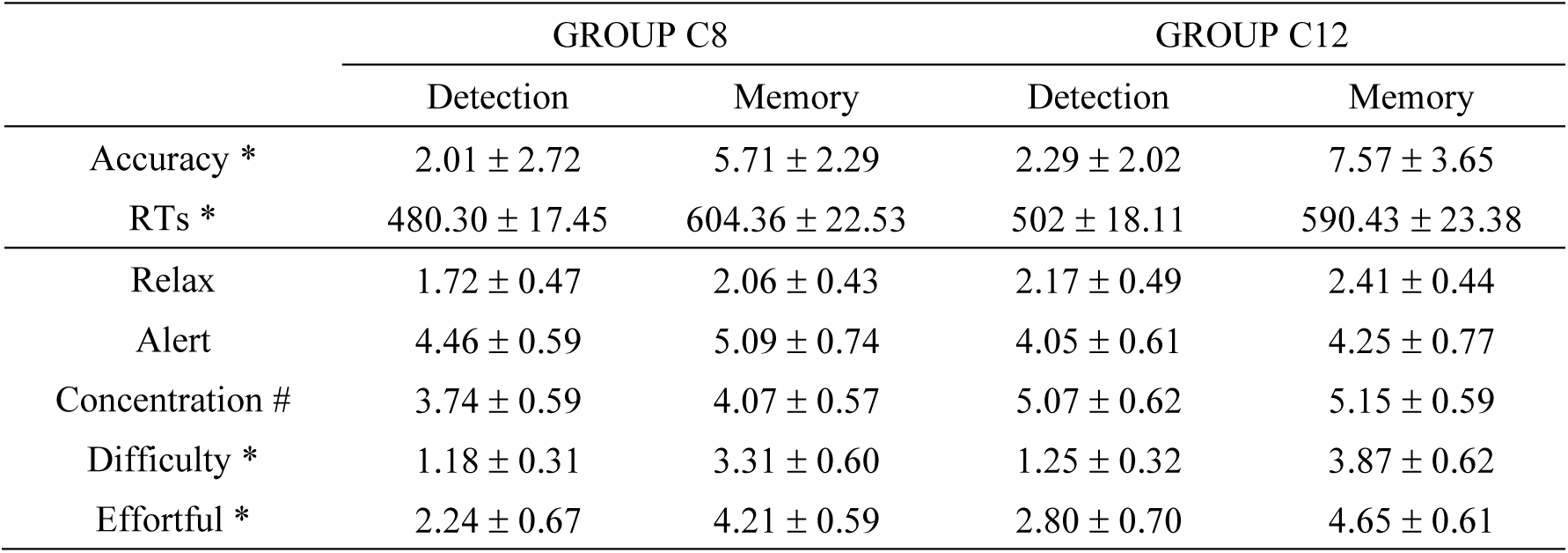
Mean (± SD) accuracy (overall error rate for all stimuli, %), reaction times (ms), and subjective ratings for the detection and memory tasks in Group C8 and Group C12. * significant main effect of Task (*p* < .05) and # borderline main effect for group (*p* = .05).

### ERP results

Figure 2a presents grand average ERPs at electrode Oz for the detection and memory tasks, together with topographic distributions at P1 and N1 peak latencies (Figure 2b), and P1–N1 difference and N1 amplitudes (Figure 2c). Scalp topographies at the P1 (98 ms) and N1 (137 ms) peak latencies (Figure 2b) confirmed that both components were maximal over occipital sites. At the N1 latency, Group C8 showed a more negative occipital distribution than Group C12 across both tasks, consistent with the significant Group effect on N1 amplitude reported below, therefore we concentrated our ERP analyses on Oz electrode traditionally used to assess early transient responses to Visually Evoked Potentials (VEPs). We also analyzed ERPs at POz, given that occipital-parietal regions are strongly associated with attention-related processes, and observed similar results to those at Oz (see Supplementary Information B).

**Figure 2.**
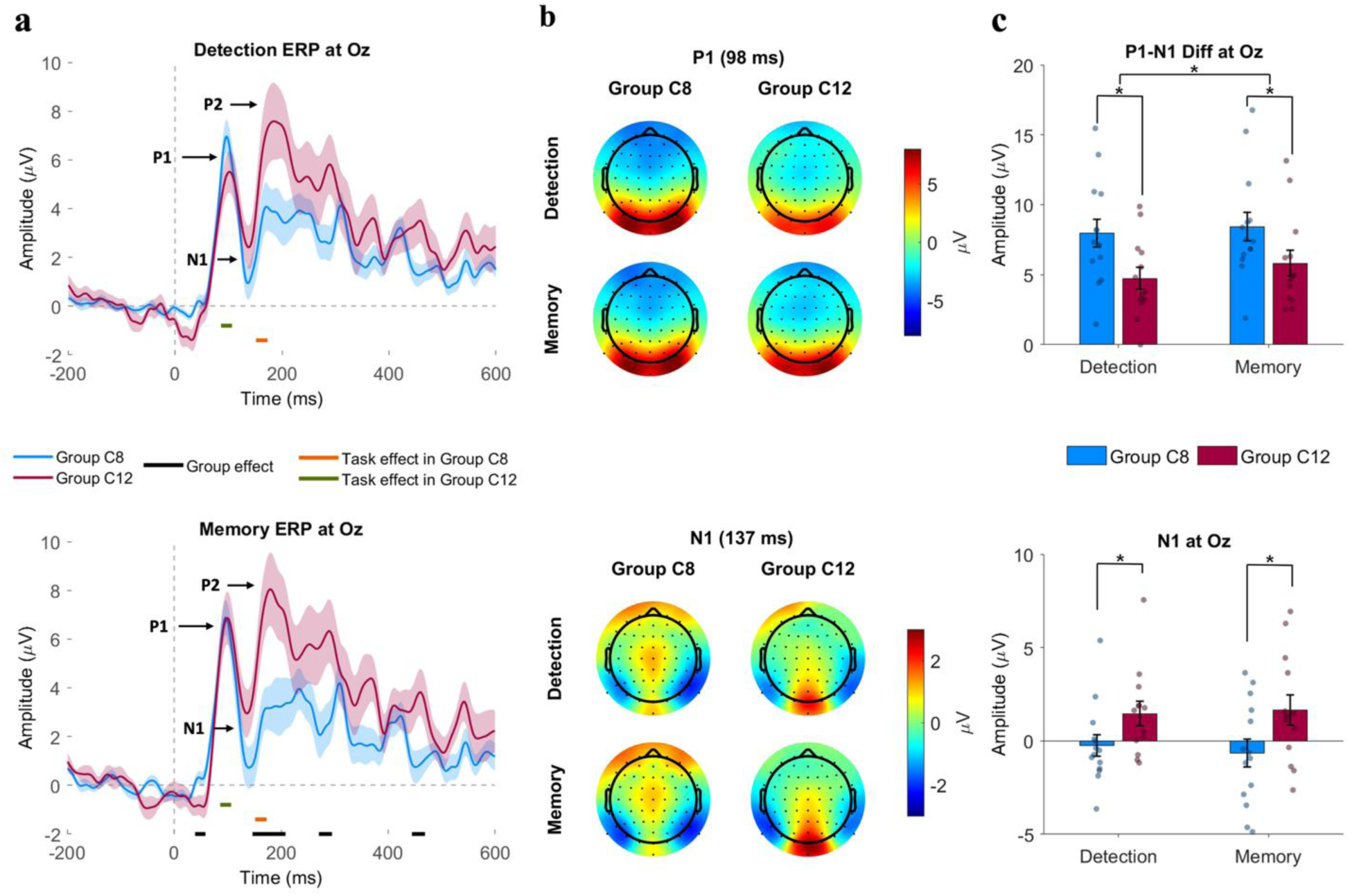
**a.** Grand average ERP waveforms at electrode Oz for detection (top) and memory (bottom) tasks. Blue lines represent Group C8 (central 8.6 Hz) and red lines represent Group C12 (central 12 Hz); shaded regions indicate ±1 SEM. Colored significance bars beneath the waveforms indicate time windows of significant point-by-point t-tests (Guthrie-corrected): black = group effect, orange = task effect in Group C8, green = task effect in Group C12. **b.** Scalp topographies at P1 (98 ms) and N1 (137 ms) peak latencies. **c.** P1–N1 peak-to-peak difference (top) and N1 amplitude (bottom) at Oz, with individual data points overlaid. Error bars represent ±1 SEM. * p < .05 after corrections applied where appropriate.

As can be observed in Figure 2a, at Oz, the latencies of the P1, N1, and P2 ERP components did not differ across the groups. Nevertheless, significant group differences were observed at multiple time points along the ERP during memory task (black horizontal bars in Figure 2a), primarily in the 150–300 ms range. We also observed differences between the memory and detection tasks: in Group C12, significant differences between the tasks emerged at approximately 50–100 ms (green bars), while task effects within Group C8 appeared around 150–200 ms (orange bars). This pattern suggests that the tagging frequency modulated the temporal window in which task-related ERP differences emerged.

For each component amplitudes (Figure 2c), no significant main effects of Task, or Task × Group interactions, were found for P1, N1, or P2 (all *p’s* > .05). However, the central tagging frequency modulated several components: N1 amplitude showed a significant main effect of Group, *F*(1, 25) = 5.11, *p* = .033, *η²* = .17, with Group C12 exhibiting more positive N1 values (*M* = 1.56 µV, *SD* = 0.64) than Group C8 (*M* = −0.45 µV, *SD* = 0.62). The Group effect on P2 amplitude approached significance, *F*(1, 25) = 4.04, *p* = .055, *η²* = .14, with Group C12 tending to show larger P2 amplitudes than Group C8 (data not shown).

To better capture early visual processing independent of baseline shifts introduced by the ongoing SSVEP oscillations, the P1–N1 peak-to-peak difference was analyzed. This measure revealed a significant main effect of Task, *F*(1, 25) = 4.31, *p* < .05, *η²* = .15, with the memory task eliciting a larger P1–N1 difference (*M* = 7.13 µV, *SD* = 0.70) than the detection task (*M* = 6.35 µV, *SD* = 0.64). The main effect of Group was also significant, *F*(1, 25) = 5.16, *p* = .03, *η²* = .17, with Group C8 showing a larger P1–N1 difference (*M* = 8.20 µV, *SD* = 0.89) than Group C12 (*M* = 5.28 µV, *SD* = 0.93). The Task × Group interaction was not significant (*p* = .436), indicating that the task-related enhancement was comparable across both groups (Figure 2c, top). These results suggest that the P1–N1 difference provides a more robust measure of task-related attentional modulation than individual peak amplitudes in the presence of concurrent SSVEP stimulation.

### SSVEP results

SSVEPs at both the tagged frequencies of 8.6 Hz and 12 Hz were observed across the occipital ROI (Figure 3a). In both groups and both tasks, the 8.6 Hz tag frequency exhibited visibly stronger SNR than the 12 Hz, as indicated by warmer colors at the 8.6 Hz horizontal line relative to 12 Hz. As expected, stronger SSVEPs were localized to posterior electrodes (see Supplementary Information C for topographic maps of raw SSVEP power). Scalp topographies (Figure 3b) confirmed that at posterior electrodes, 8.6 Hz produced broader and more intense activation than 12 Hz across all conditions. SNR time courses across the occipital ROI (Figure 3c) further illustrated group differences across the trial epoch. Within the 1000–3500 ms analysis window, Group C8 showed higher SNR than Group C12 at both tagging frequencies, while the two groups overlapped prior to 1000 ms.

**Figure 3.**
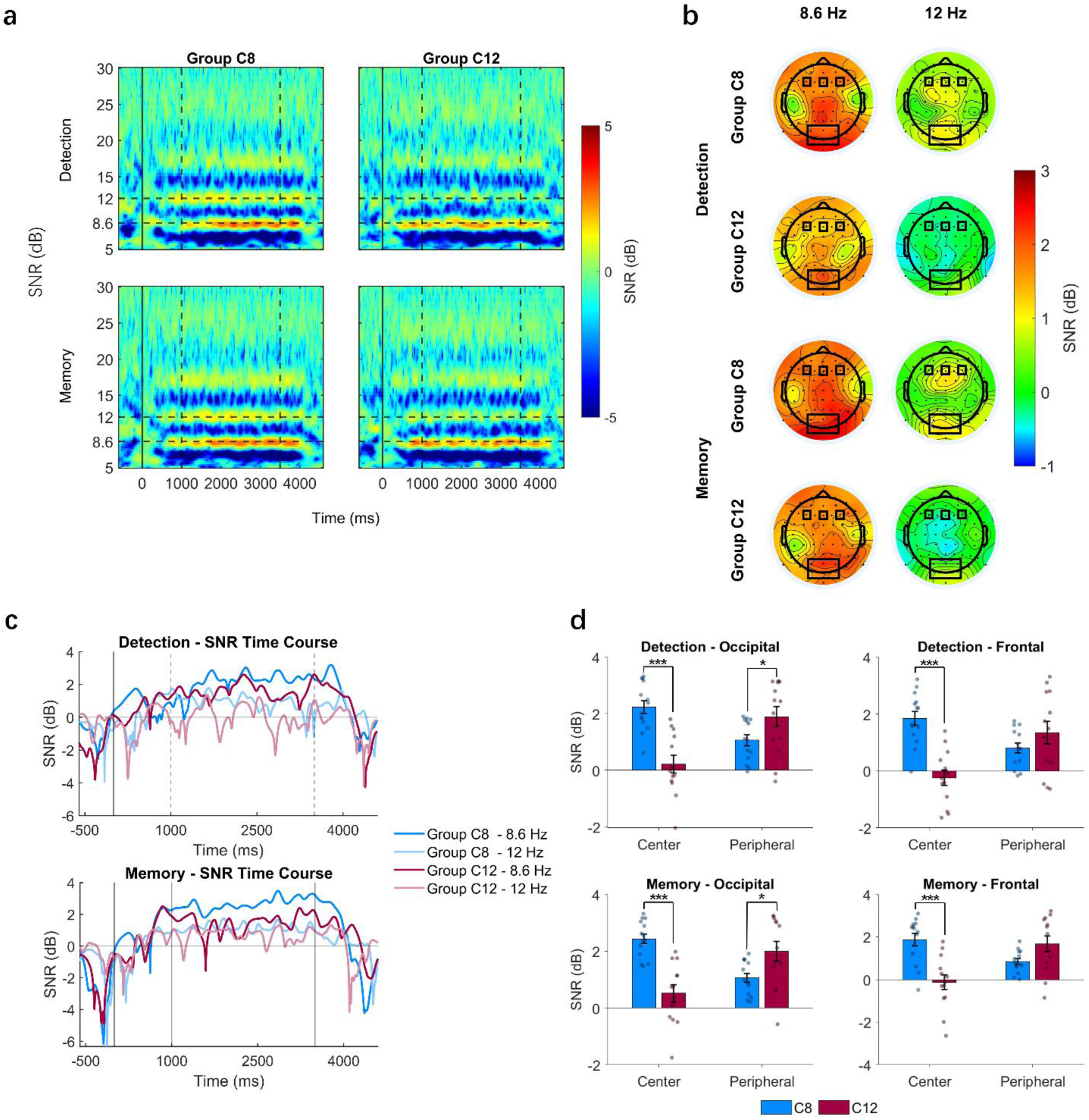
**a.** Grand average time-frequency representations of SSVEP SNR (dB) across the occipital ROI for Group C8 (left) and Group C12 (right) during the detection (top) and memory (bottom) tasks. Dashed horizontal lines indicate the 8.6 Hz and 12 Hz tagging frequencies; dashed vertical lines indicate the boundaries of the time window (1000–3500 ms) over which SSVEP SNR was averaged for topographic mapping and statistical analysis. **b.** Scalp topographies of SNR at 8.6 Hz (left) and 12 Hz (right) for each Task × Group combination. Black boxes highlight the occipital and frontal electrode cluster used for statistical analysis. **c.** SNR time courses at the 8.6 Hz and 12 Hz tagging frequencies for Group C8 and Group C12 during the detection (top) and memory (bottom) tasks at Oz electrode. **d.** Bar plots of mean SSVEP SNR across the occipital and frontal ROI at the central and peripheral tagging frequencies for the detection (top) and memory (bottom) tasks, with individual data points overlaid. Error bars represent ±1 SEM. * p < .05, *** p < .001.

A three-way mixed ANOVA with Task (Detection vs. Memory) and Location (Central vs. Peripheral) as within-subject factors and Group (C8 vs. C12) as a between-subject factor, was conducted on SSVEP SNR (dB) averaged across six occipital electrodes (O1, Oz, O2, PO3, POz, PO4) (Figure 3d, left column). Contrary to our prediction, the main effects of Task, *F*(1, 25) = 2.93, *p* = .100, *η²* = .11, and Location, *F*(1, 25) = 0.27, *p* = .611, *η²* = .01, were not significant; thus, the predicted enhancement of SSVEP magnitude during working memory relative to detection was not observed. However, there was a significant main effect of Group, *F*(1, 25) = 8.99, *p* = .006, *η²* = .26, with Group C8 showing higher overall SNR (*M* = 1.70 dB, *SD* = 0.13) than Group C12 (*M* = 1.16 dB, *SD* = 0.13).

Critically, the Location × Group interaction was highly significant, *F*(1, 25) = 23.69, *p* < .001, *η²* = .49. This interaction was predominantly driven by tagging frequency rather than stimulus location: SSVEP SNR was consistently higher at whichever location was tagged with 8.6 Hz, regardless of whether it was the attended central or the ignored peripheral location. In Group C8, where the central stimulus was tagged at 8.6 Hz, central SNR exceeded peripheral SNR (*p* = .004); in Group C12, where peripheral stimuli were tagged at 8.6 Hz, the reverse was observed (*p* < .001). Between groups, the 8.6 Hz-tagged location always produced higher SNR (central: C8 > C12, *p* < .001; peripheral: C12 > C8, *p* = .02). This pattern, together with the absence of a Location main effect, indicates that the 8.6 Hz tagging frequency dominated SSVEP signal strength, overriding any spatial attention-related enhancement that might otherwise have been expected at the attended central location (Figure 3d).

An analogous ANOVA on the frontal ROI (Fz, F3, F4; Figure 3d, right column) revealed a similar pattern. The Location × Group interaction was again highly significant, *F*(1, 25) = 18.94, *p* < .001, *η²* = .43, with a significant Group main effect, *F*(1, 25) = 10.53, *p* = .003, *η²* = .30, and no other significant effects (all *p* > .26). The two groups differed significantly for central stimuli but not for peripheral stimuli, in contrast to the pattern observed at occipital electrodes. Although the frequency-driven dominance of 8.6 Hz was not confined to occipital cortex and extended to frontal sites, unattended peripheral stimuli were less sensitive to this effect.

### The relationship between behavioural, ERP, and SSVEP measures

To further examine the relationships between SSVEPs for central and peripheral stimuli and task, subjective measures, and ERP measures, Pearson’s correlations were computed. We observed significant negative correlations between responses to centrally attended stimuli and peripheral distractors across both tasks (all *r* > .5, *p* < .01), such that increases in central SSVEP SNR were accompanied by corresponding decreases in peripheral SNR. Scatterplots for significant SSVEP SNR correlations are provided in Supplementary Information D. Consistent with the group analyses, the data showed clustering by group, with Group C8 exhibiting higher SSVEP SNRs; however, these clusters followed continuous linear trends for both central and peripheral stimuli (see Supplementary Information D). To examine whether the relationship between central and peripheral SSVEP SNR could be explained by tagging frequency, we also computed the correlation between SSVEP SNRs at 8.6 Hz and 12 Hz collapsed across all conditions. This analysis revealed a small, non-significant negative correlation (*r* = −.33, *p* = .10), however, it is important to note the possibility of a Type II error given the low effect size and sample size. Therefore, we ran partial correlations to control for the effects of tag frequency groups (see Table 2).

**Table 2.** Partial correlations controlled for frequency group (C8 and C12 groups) between occipitoparietal (OP) and frontal (F) SSVEP SNR at central attended (C) and peripheral unattended (P) stimuli in the detection and memory tasks, task performance, subjective ratings, and VEP peak component measures. \**p* < .05, ** *p* < .01 (two-tailed).

|  |  | Occipitoparietal (OP) SSVEP |  |  |  | Frontal (F) SSVEP |  |  |  |
| --- | --- | --- | --- | --- | --- | --- | --- | --- | --- |
|  |  | Detection |  | Memory |  | Detection |  | Memory |  |
|  |  | C | P | C | P | C | P | C | P |
| OP SSVEP | Detection C | -- |  |  |  |  |  |  |  |
|  | Detection P | **-.40\*** | -- |  |  |  |  |  |  |
|  | Memory C | **.65\*\*** | -.24 | -- |  |  |  |  |  |
|  | Memory P | **-.51\*\*** | **.81\*\*** | **-.42\*** | -- |  |  |  |  |
| F SSVEP | Detection C | **.83\*\*** | -.29 | **.77\*\*** | **-.46\*** | -- |  |  |  |
|  | Detection P | -.37 | **.88\*\*** | -.23 | **.70\*\*** | -.32 | -- |  |  |
|  | Memory C | **.53\*\*** | -.21 | **.77\*\*** | -.29 | **.70\*\*** | -.25 | -- |  |
|  | Memory P | **-.46\*** | **.66\*\*** | **-.41\*** | **.84\*\*** | **-.44\*** | **.77\*\*** | -.34 | -- |
| Task | Error | -- | -- | **-.51\*\*** | .23 | -- | -- | **-.50\*\*** | .14 |
|  | MRT | -.12 | -.06 | .21 | -.05 | -.01 | -.12 | .17 | -.07 |
| Subjective | Relax | .05 | -.27 | .14 | -.03 | -.16 | -.22 | .19 | -.17 |
|  | Alert | -.05 | **-.47\*** | -.06 | .04 | -.04 | -.31 | .10 | .13 |
|  | Concentration | .02 | .02 | -.19 | .08 | .04 | -.16 | -.34 | -.01 |
|  | Diff. | .12 | -.10 | .06 | -.04 | -.06 | -.24 | -.02 | -.14 |
|  | Effort | -.23 | .25 | -.11 | .23 | -.30 | .10 | -.17 | .02 |
| VEP | P1 Amp | .37 | .03 | -.09 | .11 | **.40\*** | .13 | -.11 | .32 |
|  | P1 Latency | .23 | -.28 | -.06 | -.10 | .20 | -.28 | .14 | .17 |
|  | N1 Amp | .19 | .06 | **-.39\*** | **.44\*** | .06 | .04 | -.32 | **.42\*** |
|  | N1 Latency | .28 | -.34 | .17 | -.16 | .22 | -.25 | .15 | .12 |
|  | P2 Amp | -.10 | .21 | -.35 | .28 | -.11 | .23 | -.26 | .33 |
|  | P2 Latency | .03 | -.14 | -.03 | -.28 | .03 | -.10 | -.05 | -.12 |
|  | P1–N1 Diff | .20 | -.27 | .22 | -.24 | .33 | -.13 | .15 | -.03 |

Occipitoparietal and frontal SSVEP SNR was highly consistent with each other, and between the detection and memory tasks, with strong positive correlations between detection and memory conditions for both central and peripheral locations (*r* > .65, *p* < .01 for all, see Table 2). Occipitoparietal central and peripheral SSVEP SNR were negatively correlated in both the detection (*r* = −.40, p < .05) and memory (*r* = −.42, p < .05) tasks after controlling for the effects of tag frequency groups. However, these negative correlations were not significant for frontal SSVEP SNR in either task (see Table 2). Together, these results suggest the negative correlation between SSVEP SNR for centrally attended and peripheral distraction may reflect differences in attention allocation and cannot be fully accounted for by tag frequency.

For ERP measures, only P1 amplitude significantly correlated with frontal SSVEP SNR for central attended stimuli in the detection task after controlling for group (partial *r* = .40, p < .05). For the memory task, N1 amplitude significantly correlated with occipitoparietal SSVEP SNR for both centrally attended (partial *r* = −.39, p < .05) and peripheral distractor (partial *r* = .44, p < .05) stimuli. Interestingly, these opposing directions are consistent with the negative correlation between central and peripheral SSVEP SNR (see Table 2).

Behavioural measures and subjective ratings also showed significant associations with SSVEP SNR. For occipitoparietal SSVEP, alertness was negatively correlated with peripheral SSVEP SNR during the detection task (partial *r* = −.47, *p* < .05); The more alert participants showed lower SSVEP SNR for the periphery stimuli in the detection task, consistent with more effective distractor suppression. For the memory task, higher central SSVEP SNR was associated with fewer errors at both frontal (partial *r* = −.50, *p* < .01) and occipitoparietal sites (partial *r* = −.51, *p* < .01), suggesting that stronger neural entrainment to the central stimulus supported working memory performance. Accuracy on the detection task was very high and violated the assumption of normality, therefore correlations were not run. No significant correlations were found between SSVEP SNR and reaction times in either task. To further explore this relationship, we ran an exploratory Stepwise Regression to ascertain the EEG measures able to predict accuracy of the memory task; central and peripheral for both occipitoparietal and frontal SSVEP SNR, P1, N1, P2 amplitudes, differences between P1-N1 amplitudes were entered as predictors for error rate on the memory task (i.e., overall accuracy). Group was also entered as a control. Only occipitoparietal SSVEP SNR for central stimuli (part-partial *r* = −.48, *p* = .008) was identified as a significant predictor of accuracy on the memory task, *R* = .57, *R^2^*= .33, *F*(1,25) = 5.84, *p* = .009, accounting for 23% of unique variance.

## Discussion

The present study used a dual frequency-tagging paradigm with 8.6 Hz and 12 Hz tags to examine the sensitivity of SSVEP and ERP measures to detection versus working memory task demands with distractors. SSVEP signal strength was strongly determined by tagging frequency, with 8.6 Hz consistently producing higher SNR than 12 Hz regardless of spatial position, a pattern that extended from occipital to frontal sites. Although time-averaged SSVEP SNR was not significantly different between the two tasks at a group level contrary to our initial hypothesis, SSVEP SNR for centrally attended stimuli and unattended peripheral distractors were negatively correlated even after controlling for the effects of tag-frequency group. Time-averaged SSVEPs for central attended stimuli also significantly predicted task performance on the memory task. Task-related modulation was also observed in the ERP component (P1–N1 peak-to-peak difference), which was significantly larger during working memory than detection. Point-by-point comparisons further revealed task-dependent ERP modulations whose temporal profile varied with tagging frequency.

We selected 8.6 Hz and 12 Hz to remain within the alpha band range commonly used in SSVEP attention studies [7, 8, 18]. While early frequency-tagging studies treated tagging frequency as a neutral means of separating stimuli [7], subsequent work has demonstrated that responses depend on the chosen frequency, particularly within the alpha range [8], and more broadly on the neural networks entrained by different frequencies [18]. Consistent with these studies, we observed higher SSVEP SNR for 8.6 Hz than 12 Hz regardless of stimulus location. This asymmetry may reflect the resonance peak of visual cortex in the lower alpha band range [29, 30], and aligns with the functional dissociation within the alpha band whereby lower alpha oscillations (∼8–10 Hz) are more closely linked to general attentional processes such as alertness and arousal, whereas upper alpha activity (∼10–12 Hz) predominantly reflects task-specific semantic processing [31, 32]. The proximity of the 8.6 Hz tag to the lower alpha range may thus have preferentially engaged attention-related neural mechanisms, consistent with the frequency-dependent network recruitment reported by Ding et al. [18]. Interestingly, this neural asymmetry was paralleled by subjective ratings: participants in Group C8 reported the task as marginally easier to concentrate on, suggesting that subjective experience may also be sensitive to tagging frequency. Nevertheless, significant relationships were observed between SSVEP responses to centrally attended and peripheral stimuli that could not be fully accounted for by tagging frequency. Specifically, and consistent with prior research, increases in SSVEP amplitude for centrally attended stimuli were associated with decreases in SSVEP amplitude for unattended peripheral stimuli. In addition, frontal sites appeared to show reduced sensitivity to the 8.6 Hz tagging frequency assigned to the peripheral unattended distractors. Future studies should systematically vary tagging frequencies within groups and across lower and upper alpha sub-bands to disentangle resonance-driven and attention-driven SSVEP modulations.

The tagging frequency also influenced early ERP, components prior to the onset of SSVEPs, with important interpretive implications. The N1 component showed a significant Group effect, with more negative N1 values in Group C8, yet peak latencies did not differ across groups. The N1 group effect likely reflects a direct influence of tagging frequency on the early transient visual response. At the N1 latency (∼137 ms), only one to two flicker cycles would have elapsed in our study, and the steady-state response would likely not have yet been established [20, 33], indicating that the observed amplitude difference originated from the transient onset response rather than from superimposition of an ongoing oscillation. The 8.6 Hz tag, falling within the lower alpha range, may have produced larger early cortical responses through closer alignment with the natural resonance frequency of visual cortex [29, 30]. Consistent with this, the P1–N1 peak-to-peak difference which controls for baseline shifts, retained a significant Group effect (C8 > C12), confirming a genuine enhancement of the transient response evoked by the lower tagging frequency [34].

Critically, the choice of tagging frequency shaped not only signal characteristics but also the temporal window in which task specific effects became measurable. Point-by-point task comparisons within each group revealed that task-related ERP differences emerged earlier in Group C12 (approximately 50–100 ms) and later in Group C8 (approximately 150–200 ms). While previous studies have examined how attention modulates SSVEPs at different frequencies within the alpha band [8, 18], these findings address a complementary question: the tagging frequency itself, as a methodological parameter, systematically influences SSVEP signal strength, ERP component morphology, and the detectability of task specific effects within paradigms.

Working memory has been associated with SSVEP modulations at frontal sites [26–28]. In this study, however, SSVEP SNR did not significantly differ between the detection and memory tasks at frontal or occipitoparietal sites at a within group level despite observed behavioral differences in accuracy and motor speed. Both tasks required sustained spatial attention to the central stimulus and employed identical visual displays, so the sustained sensory processing indexed by occipitoparietal SSVEPs was likely equivalent across conditions. However, distractor effects on attention are known to depend on both feature similarity [12] and task load [14]. In this context, the combined cognitive demands of the tasks, together with the similarity between attended and distractor features, may not have been sufficiently distinct to elicit measurable differences in sustained SSVEP responses. Increasing task demands, for example through a higher memory load such as a 2-back paradigm, may be necessary to reveal task- and distractor-relevance-related modulation of sustained SSVEP activity.

Although the main grouped SSVEP analyses did not reveal significant task effects, correlational analyses revealed systematic links between SSVEPs for behavior, neural responses, and subjective state. Several patterns were consistent with altered allocations of attentional resources; increases in SSVEPs for centrally attended stimuli were significantly associated with decreases in SSVEP SNR for peripheral distractors [5, 35]. In addition, during the memory task, stronger central SSVEP SNR at both frontal and occipitoparietal sites was related to lower error rates, though only occipitoparietal SSVEP SNR was identified as a unique predictor of accuracy on the memory task. We also observed associations between SSVEP SNR and early transient P1 and N1 ERP amplitudes, indicating shared attentional gain across steady-state and early transient visual mechanisms. Complementing these neural measures, lower self-reported alertness was associated with higher peripheral occipitoparietal SSVEP SNR during detection, suggesting that reduced alertness weakened suppression of peripheral distractors as predicted [35, 36].

Several methodological limitations should be acknowledged. The tag frequency-to-location mapping was assigned between subjects, so the observed Group effects may partly reflect pre-existing individual differences; a within-subject design would provide a more controlled comparison for assessing the effects of tag frequency (8.6 Hz vs. 12 Hz) on attention. As discussed above, the proximity of 8.6 Hz to the lower alpha band resulted in a frequency-dependent asymmetry that dominated the SSVEP results. This confound may have been further compounded by individual differences in alpha peak frequency, which modulates the interaction between flicker-driven and endogenous oscillations [34], but was not measured in the present study. Future work should incorporate individual alpha peak frequency as a covariate and consider tagging frequencies outside the alpha range to minimize resonance-related confounds.

In conclusion, contrary to our prediction, SSVEP SNR did not differ between detection and 1-back working memory tasks at a group level, but was strongly determined by tagging frequency, with 8.6 Hz consistently eliciting stronger responses than 12 Hz regardless of location or task, a pattern that extended to frontal sites. Nevertheless, central SSVEP SNR predicted working memory accuracy, and the P1–N1 peak-to-peak difference was significantly larger during the memory task, suggesting that transient ERP measures may capture task demands that sustained SSVEP measures do not. A mere 3.4 Hz difference in tagging frequency was sufficient to alter SSVEP signal strength, shift ERP component amplitudes, and change the temporal window in which cognitive effects emerged, highlighting that tagging frequency should be treated as an active experimental variable rather than a neutral methodological choice.

## Acknowledgements

We thank Professor Neil Levy (N.L.) for his support and advice on study design. Thank you to the Florey Institute of Neuroscience and Mental Health, Melbourne, Australia for supporting data collection for this study. This work was financially supported by a grant from the John Templeton Foundation awarded to N.L. and R.H., Florey Institute of Neuroscience and Mental Health, Melbourne, Australia.

## Author contributions

Jihan Yang - analysis and write-up of first draft, review and approval of manuscript.

Olivia Carter - oversight and supervision, experimental design, set-up, review and approval of manuscript.

Mohit N Shivdasani - supervision, analysis, write-up and final review.

David Grayden - experimental design, set-up, analysis, review and approval of manuscript.

Rob Hester - experimental design, set-up, review and approval of manuscript.

Ayla Barutchu - supervision, experimental design, set-up, data collection and collation, analysis, write-up, review and approval of manuscript.

## Data Availability Statement

Data is available upon request from the corresponding author.

