## Supplementary Information for "When Tagging Frequency Matters to Attention: Effects on SSVEPs, ERPs, and Cognitive Processing"

### Supplementary Information A

**Grand-averaged ERP waveforms at Oz across the entire trial epoch**

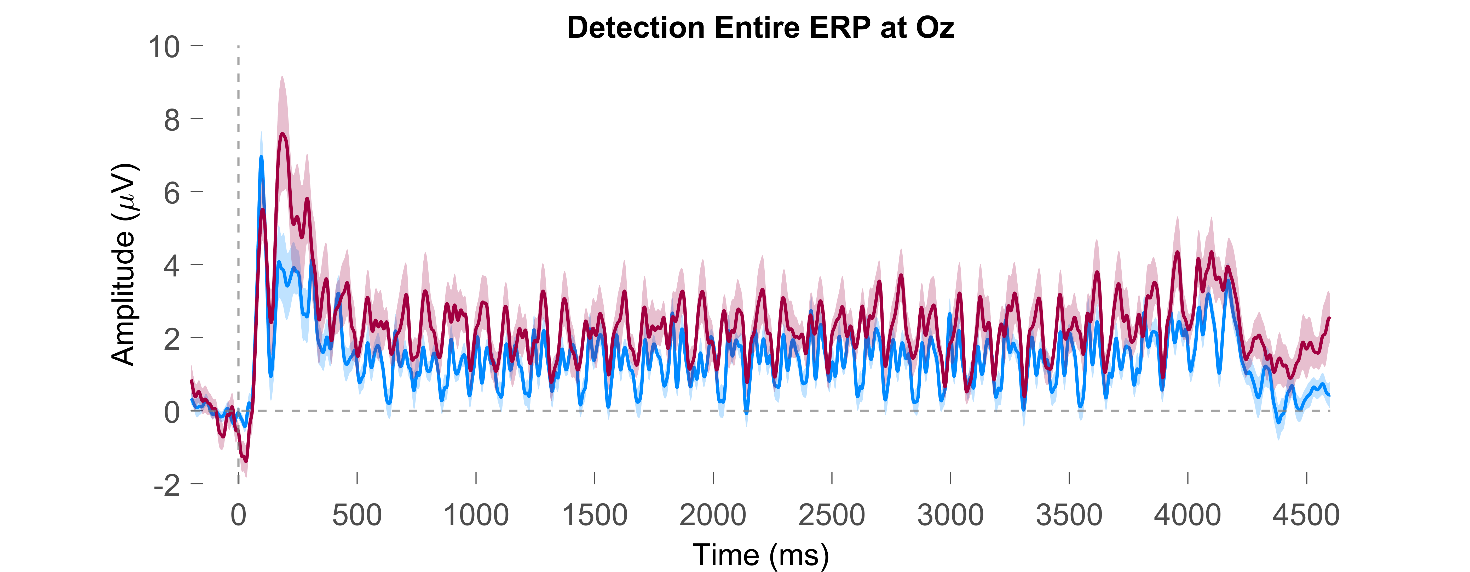

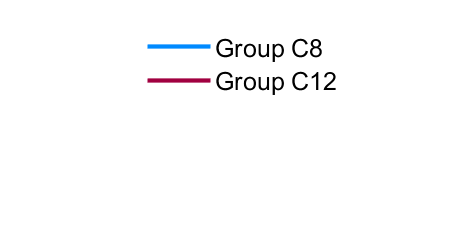

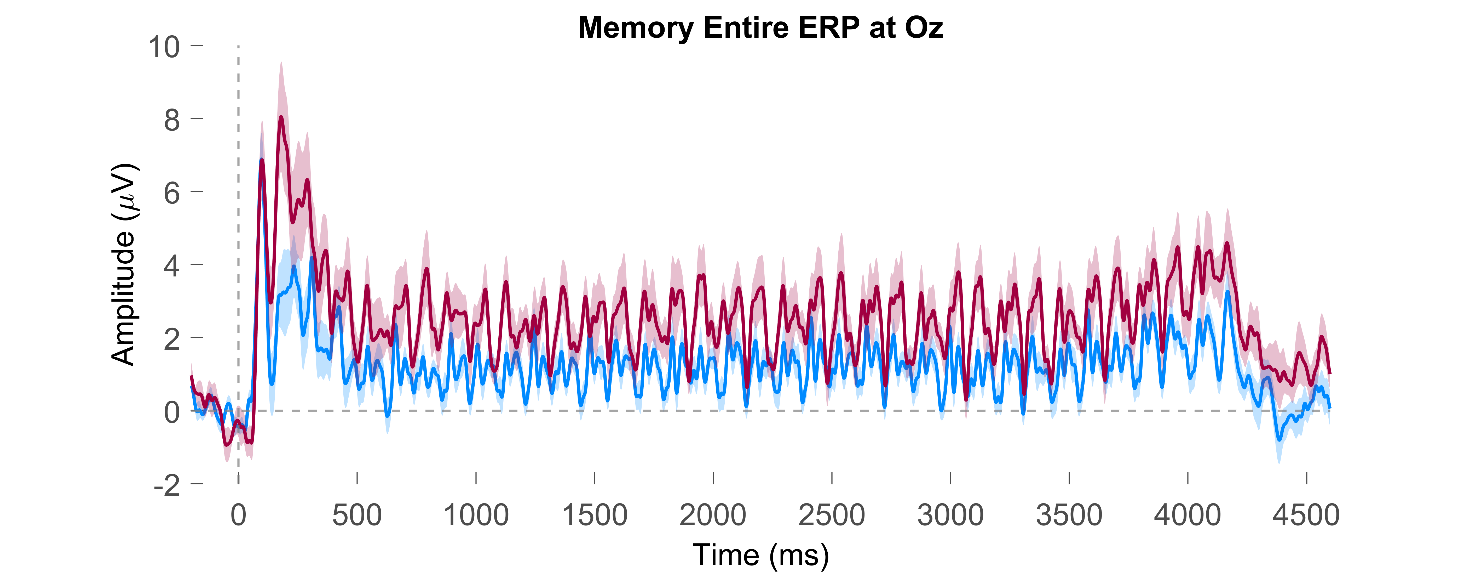

*Figure SA 1*. Grand-averaged ERP waveforms at electrode Oz for the detection (top) and memory (bottom) tasks across the entire trial epoch (-200–4600 ms). Blue and red lines represent Group C8 (central 8.6 Hz) and Group C12 (central 12 Hz), respectively; shaded regions indicate ±1 *SEM*. The dashed vertical line indicates stimulus onset.

### Supplementary Information B

**Grand-averaged ERP waveforms at POz**

a

| 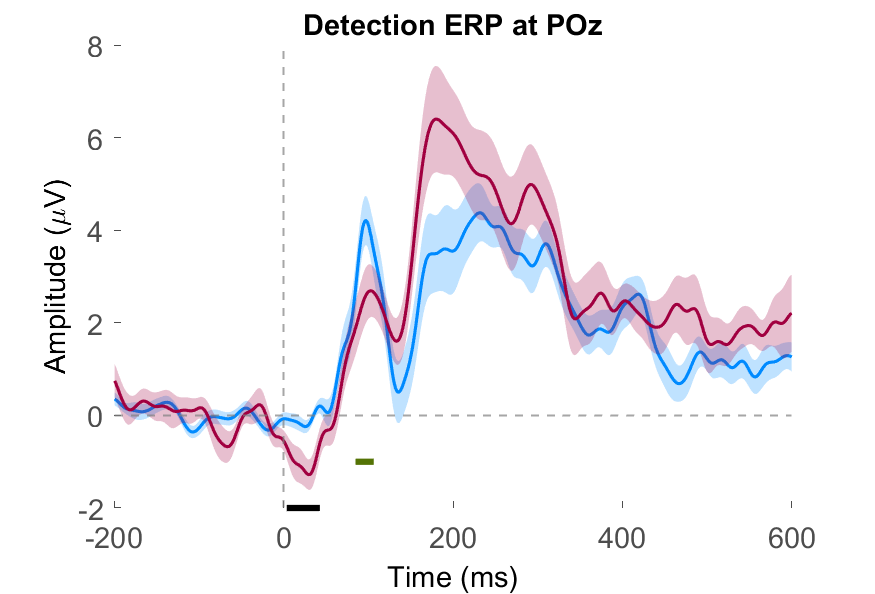 | 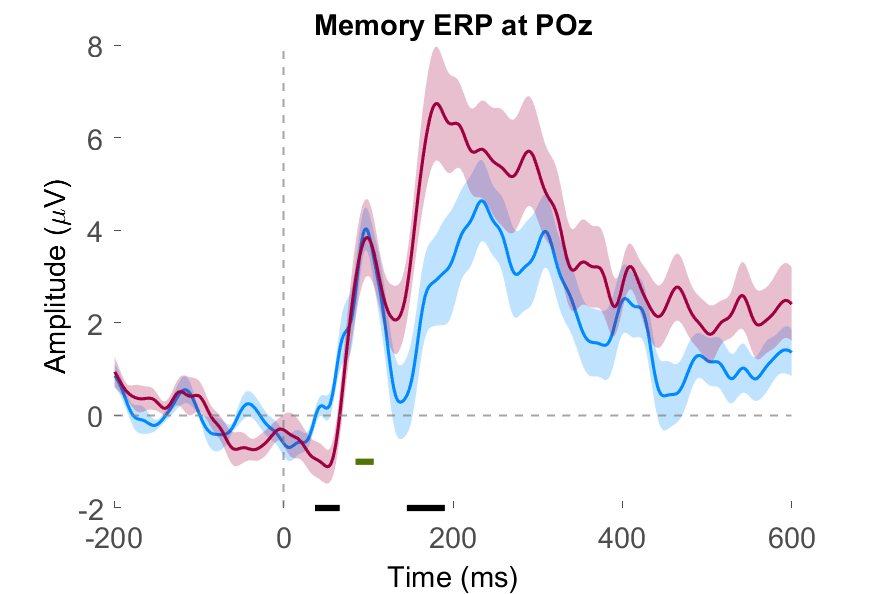 |
| --- | --- |

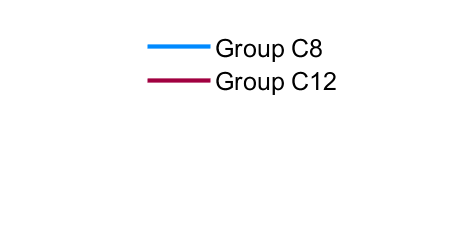

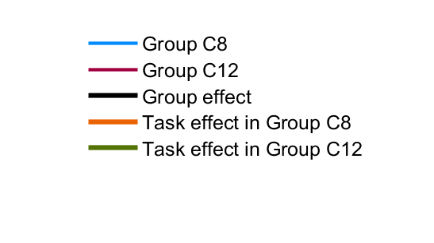

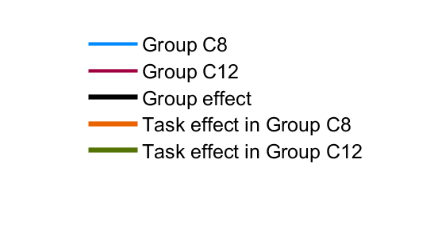

b

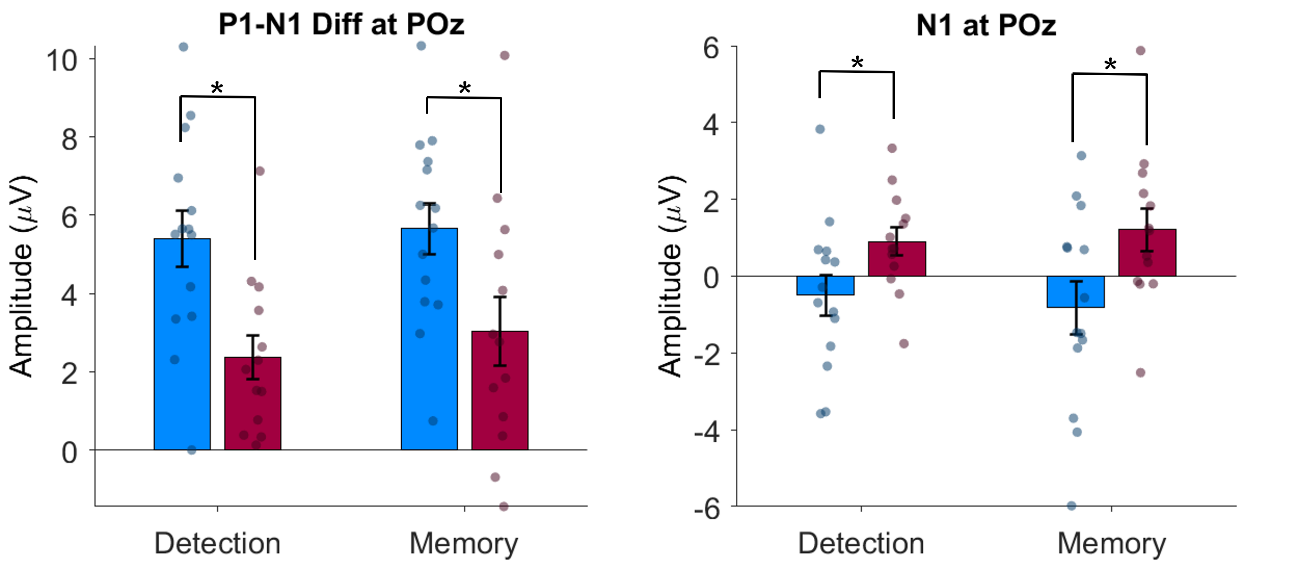

*Figure SB 1*. **a**) Grand-averaged ERP waveforms at electrode POz for the detection (top left) and memory (top right) tasks over the first 600 ms post-stimulus. Blue and red lines represent Group C8 and Group C12, respectively; shaded regions indicate ±1 *SEM*. The black horizontal bar indicates time points where the group effect reached significance and green horizontal bar indicates task effect in Group C12 (Guthrie-corrected point-by-point paired-samples t-tests, α = .05, minimum 20 consecutive ms). **b**) Mean P1–N1 difference and N1 amplitude at POz. Peak-to-peak difference (left) and N1 amplitude (right) at electrode POz for the detection and memory tasks, with individual data points overlaid. Blue and red bars represent Group C8 and Group C12, respectively. Error bars represent ±1 *SEM*. * p < .05 after corrections applied where appropriate.

### Supplementary Information C

**Time-frequency representations of raw SSVEP power at Oz.**

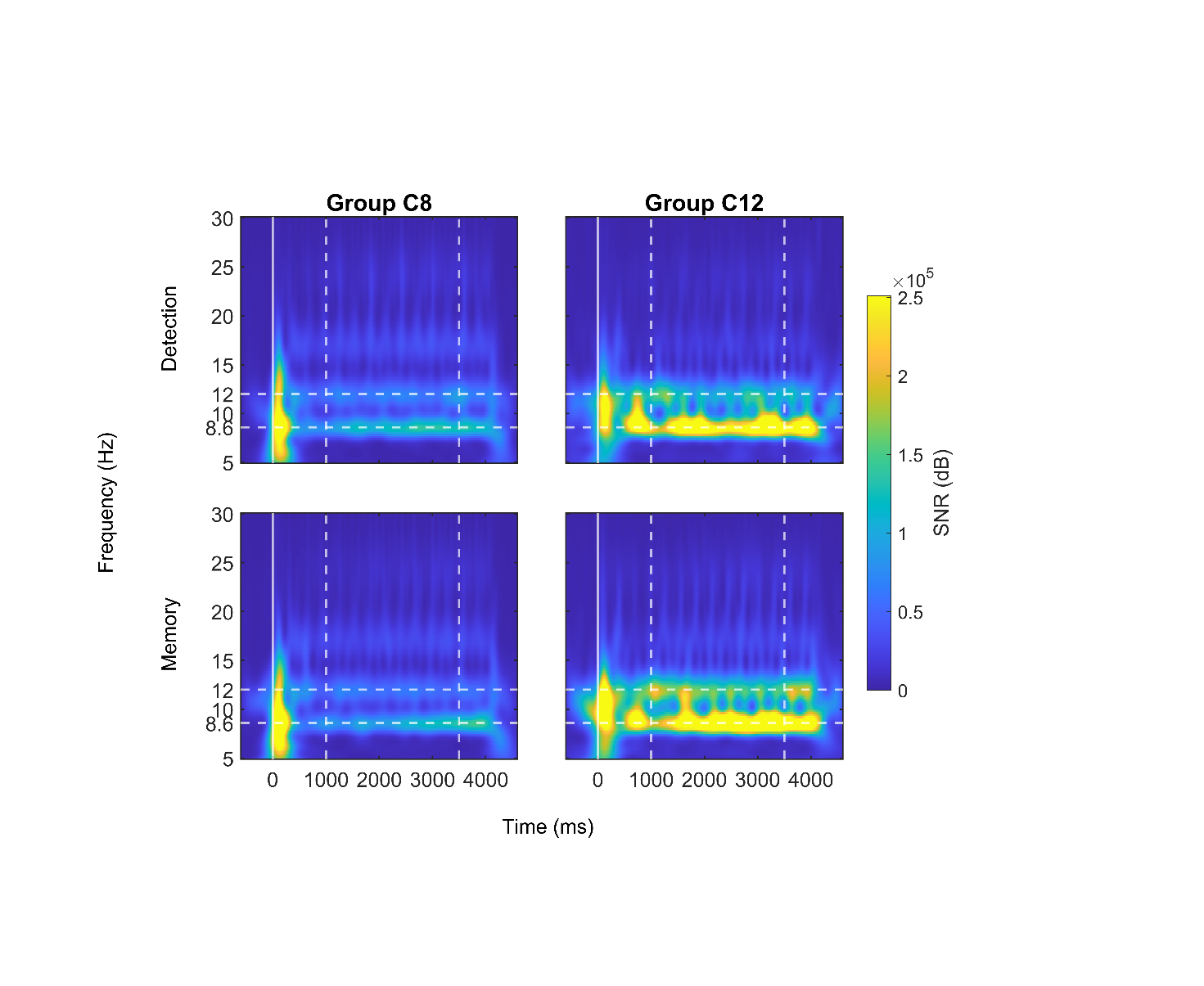

*Figure SC 1*. Time-frequency spectrograms of raw SSVEP evoked power (μV²) at electrode Oz for Group C8 (left) and Group C12 (right) during the detection (top) and memory (bottom) tasks. Dashed horizontal lines indicate the 8.6 Hz and 12 Hz tagging frequencies; dashed vertical lines indicate the boundaries of the time window (1000–3500 ms) over which SSVEP SNR was averaged for topographic mapping and statistical analysis.

**Scalp topographies of raw SSVEP evoked power**

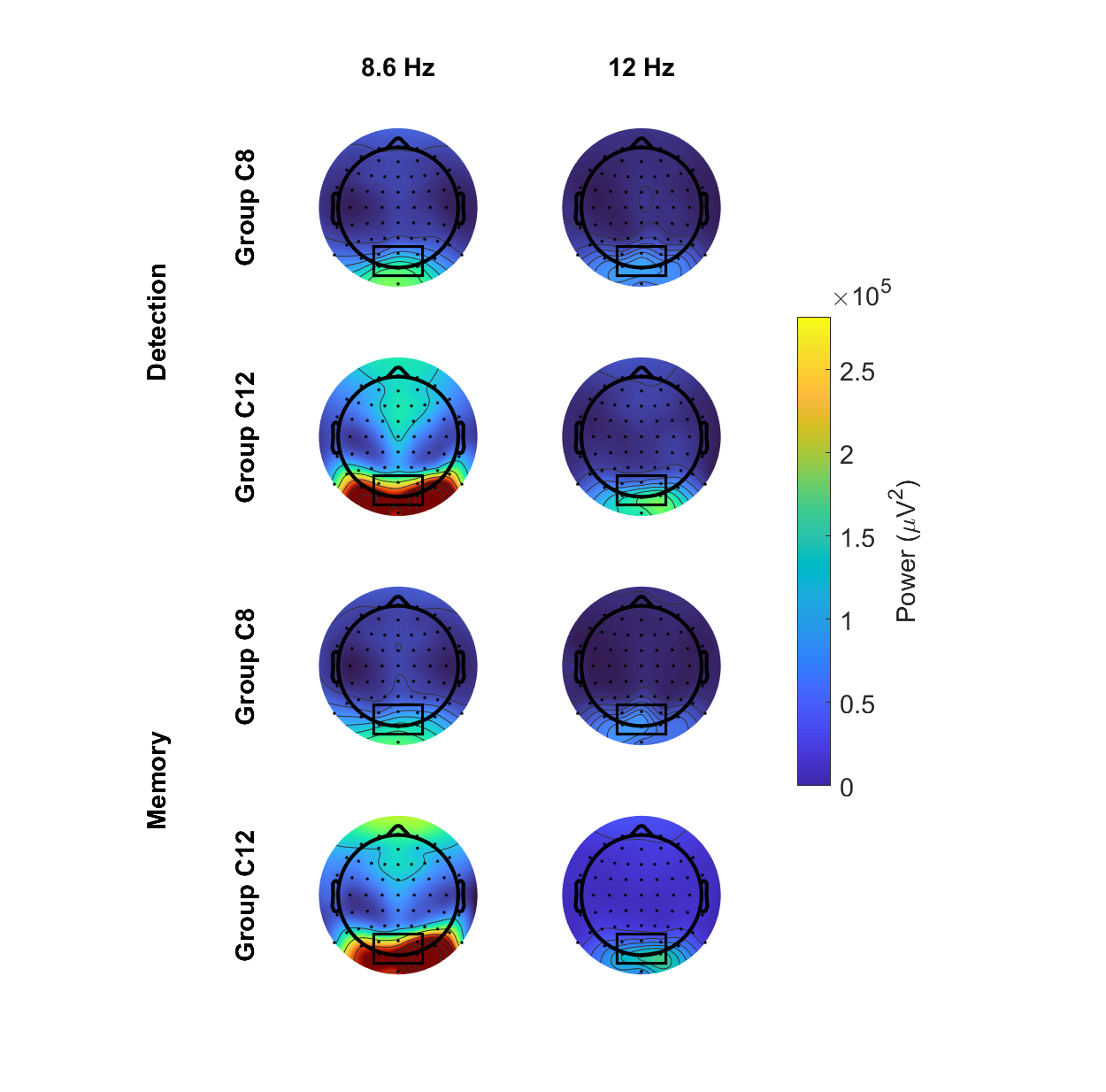

*Figure SC 2*. Scalp topographies of raw SSVEP evoked power (μV²) at 8.6 Hz (left) and 12 Hz (right) for the detection (top two rows) and memory (bottom two rows) tasks, shown separately for Group C8 and Group C12. The color scale is shared across all panels.

### Supplementary Information D

**Scatter plots for Significant Pearson Correlations**

| 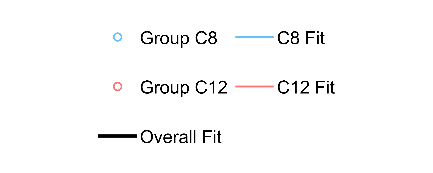 | | | |
| --- | --- | --- | --- |
| 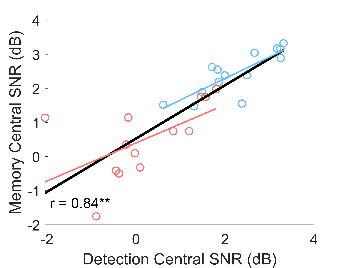  a | 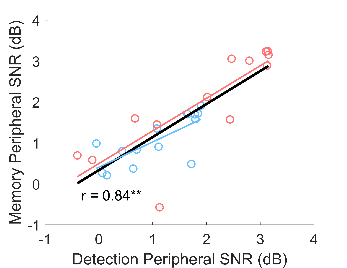  b | 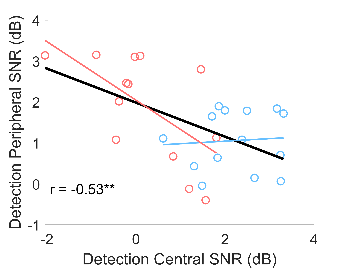  c | 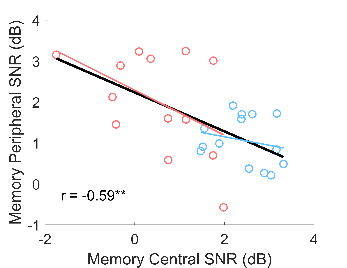  d |

*Figure SD 1*. Scatter plots of significant Pearson correlations between occipitoparietal SSVEP SNR measures across conditions: **a)** Detection Central vs Memory Central SNR (partial *r* = .65, *p* < .01), **b)** Detection Peripheral vs Memory Peripheral SNR (partial *r* = .81, *p* < .01), **c)** Detection Central vs Detection Peripheral SNR (partial *r* = -.40, *p* < .01), **d)** Memory Central vs Memory Peripheral SNR (partial *r* = -.42, *p* < .05). Note that although the groups are clustering, they are nevertheless following the same linear trend, and partial correlations remain significant after controlling for the effects of group. Figures depict significant Pearson’s correlations (** = *p* < .01).

a

| 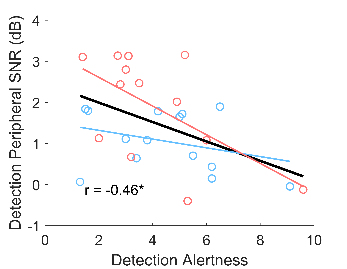  b | 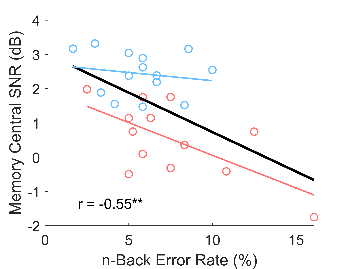  c | 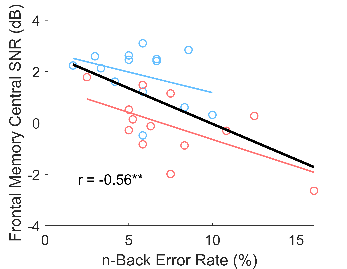 |
| --- | --- | --- |

*Figure SD 2.* Scatter plots of significant Pearson correlations between SSVEP SNR and behavioral measures. **a)** Occipitoparietal Detection Peripheral SNR vs Detection Alertness (partial *r* = -.47, *p* < .05), **b)** Occipitoparietal Memory Central SNR vs n-Back Error Rate (partial *r* = -.51, *p* < .01), **c)** n-Back Error rate vs Frontal Memory Central SNR (partial *r* = -.50*, p* < .01). Partial correlations controlling for the effects of group.
